# Genome-Wide Identification of HES1 Target Genes Uncover Novel Roles for HES1 in Pancreatic Development

**DOI:** 10.1101/335869

**Authors:** Kristian Honnens de Lichtenberg, Nina Funa, Nikolina Nakic, Jorge Ferrer, Zengrong Zhu, Danwei Huangfu, Palle Serup

**Affiliations:** Novo Nordisk Foundation Center for Stem Cell Biology (Danstem), University of Copenhagen, Imperial College London, UK; Section on Epigenetics and Disease, Imperial College London, UK; Center for Stem Cell Biology, Memorial Sloan Kettering Cancer Center, NEW YORK; Current Address: GSK, Stevenage, UK

## Abstract

Notch signalling and the downstream effector HES1 is required for multiple pancreatic cell fate choices during development, but the direct target genes remain poorly characterised. Here we identify direct HES1 target genes on a genome-wide scale using ChIP-seq and RNA-seq analyses combined with human embryonic stem cell (hESC) directed differentiation of CRISPR/Cas9-generated *HES1*^*-/-*^ mutant hESC lines. We found that HES1 binds to a distinct set of endocrine-specific genes, a set of genes encoding basic Helix-Loop-Helix (bHLH) proteins not normally expressed in the pancreas, genes in the Notch pathway, and the known HES1 target NEUROG3. RNA-seq analysis of wild type, *HES1*^-/-^, *NEUROG3*^-/-^, and *HES1*^-/-^*NEUROG*3^-/-^ mutant hESC lines allowed us to uncover NEUROG3-independent, direct HES1 target genes. Among the HES1 bound genes that were derepressed in *HES1*^-/-^*NEUROG3*^-/-^ cells compared to *NEUROG3*^-/-^ cells, we found members of the endocrine-specific gene set, the Notch pathway genes *DLL1*, *DLL4*, and *HEY1*, as well as the non-pancreatic bHLH genes *ASCL1* and *ATOH1*. We also found a large number of transcripts specific to the intestinal secretory lineage to be increased in *HES1*^-/-^*NEUROG3*^-/-^ cells. Together, our data reveal that HES1 employs a multi-layered control of endocrine differentiation, controls Notch ligand expression independent of NEUROG3, and prevents initiation of ectopic intestinal transcriptional programmes in pancreas progenitors.

## Introduction

Stem cell-based regenerative therapy for Type I Diabetes has the potential to prevent secondary complications and relieve patients from cumbersome blood glucose measurements multiple times per day by providing transplantable, in vitro differentiated insulin-producing β-cells. Recent years has brought great advances in protocols to produce β-like-cells from human embryonic stem cells (hESCs), largely based on knowledge from pancreas development in the mouse and other model organisms. *In vitro* differentiation of hESCs into β-like-cells also presents an opportunity to study a simplified, scalable and accessible model of human pancreas development.

The mouse pancreas epithelium consists of three tissue types, enzyme producing acinar cells, hormone producing endocrine cells of the islet of Langerhans and duct cells lining the ductal tree that connects to the small intestine. These parenchymal cells all stem from a common multipotent progenitor cell (MPC) population, marked by co-expression of a signature set of transcription factors; Pdx1, Nkx6-1, Ptf1a and Sox9. Later, the MPCs segregate into Ptf1a+ tip-and Sox9+Nkx6-1+ trunk progenitors under the influence of a mutually repressive interaction between Ptf1a and Nkx6-1 and instructed by Notch signalling (Afelik et al., 2012; Horn et al., 2012; Shih et al., 2012). Endocrine cells arise from an intermediary precursor stage marked by Ngn3 expression (Apelqvist et al., 1999; Gradwohl et al., 2000; Gu et al., 2002; Jensen et al., 2000a; Schwitzgebel et al., 2000). The first endocrine cells emerge from MPCs through and are mainly α-and ε-cells, while β-, δ-, and γ-cells arise later from the trunk epithelium (Heller et al., 2004; Johansson et al., 2007; Prado et al., 2004). We and others have shown that loss of Hes1 as well as other Notch pathway genes leads to excess endocrine differentiation (Apelqvist et al., 1999; Fukuda et al., 2006; Jensen et al., 2000b; Sumazaki et al., 2004). Thus, formation of endocrine cells is negatively regulated by Notch such that low Notch activity and thereby low Hes1 levels relieve *Neurog3* repression and allow high levels Neurog3 protein and thereby commitment to the endocrine lineage, Neurog3 is expressed only transiently (Bankaitis et al., 2015; Gradwohl et al., 2000; Gu et al., 2002; Jensen et al., 2000a) while located in the progenitor epithelium after which the differentiating endocrine cells, now expressing Neurog3 target genes such as Neurod1 and Insm1, delaminate from the epithelium and coalesce into the islets of Langerhans as single hormone producing α-, β-, γ-, δ-and ε-cells (Gasa et al., 2004; Gradwohl et al., 2000; Jensen et al., 2000b; Schwitzgebel et al., 2000).

Notch signalling thus plays a pivotal role in pancreas cell fate choice, but little is known about how Notch and its main effector Hes1 regulates pancreas development on a molecular level. Generally, Notch signalling is activated by Delta/Jagged-family ligands that bind Notch receptors on a neighbouring cell inducing a series of proteolytic cleavages the last of which is by the ?-secretase complex. This leads to translocation of the Notch intracellular domain to the nucleus, where it acts in complex with Rbpj and Maml1 to activate transcription (Bray, 2006). Prominently among the target genes activated by Notch in the pancreas is *Hes1*, encoding a bHLH-protein that itself binds to specific regulatory sites on DNA and recruits Groucho/TLE proteins to form a repressive complex (Kageyama et al., 2007). Only a few Hes1 target genes are described in the pancreas, including *Neurog3* and *Cdkn1c*, encoding p57^Kip2^(Georgia et al., 2006; Lee et al., 2002). Notably, in several tissues including presomitic mesoderm and neural progenitors, but not yet described in the pancreas, Hes1 binding to its own promoter results in auto-repression and oscillatory expression of the protein as well as of some of its target genes (Hirata et al., 2002; Kobayashi et al., 2009a; Sasai et al., 1992; Takebayashi et al., 1994). Other examples of negative feedback in the Notch pathway includes lateral inhibition with its associated suppression of ligand expression in cells that receive Notch signalling, typically as a consequence of Hes-mediated repression of bHLH family transcriptional activators that activate ligand expression in the absence of Notch (Bray, 2006).

Recent work demonstrates that many of the same transcription factors are found in the human embryonic pancreas, and in much the same patterns (Jennings et al., 2017; 2013). However, since the fetal development of humans compared to mice is much longer and due to the scarcity of human fetal material from early stages it is unclear whether there is a clear equivalent to the primary/secondary transition stages seen in rodents and whether Notch signalling plays a similar role in human development as in mouse. Nevertheless, most hESC protocols aimed at generating β-like-cells successfully employ γ-secretase inhibitors to induce endocrine differentiation indicating that Notch signalling is active in hESC-derived pancreas progenitors. Moreover, recent work from one of us revealed that CRISPR/Cas9-mediated *HES1* null mutations in hESCs cause increased formation of C-peptide^+^ cells (Zhu et al., 2016), bolstering the case for Notch-mediated regulation of pancreatic endocrine development in humans also.

Here we take advantage of the ease at which pancreatic progenitors can be produced from hESCs to define direct HES1 target genes on a genome-wide scale and thereby uncover novel functions for HES1 in pancreatic development. We identify genomic HES1 binding sites by chromatin immunoprecipitation followed by deep sequencing (ChIP-seq) and compare bound genes to genes that are deregulated by CRISPR/Cas9-induced loss of *HES1*. Additional deletion of HES1 on a NEUROG3-null background enabled us to assess genes deregulated by loss of *HES1* independently of the endocrine programme initiated by NEUROG3. Thereby we uncovered a subset of endocrine-specific transcripts that are regulated in opposite directions by NEUROG3 and HES1 and show derepression in *HES1*^-/-^*NEUROG3*^-/-^ double-null cells. We also unravel a direct repression of Notch ligand genes *DLL1* and *DLL4* by *HES1*. A number of HES1 bound distal regulatory elements map to known FOXA2 binding sites suggesting co-occupancy with FOXA2. Remarkably, these targets include lineage specific genes that are not expressed in the pancreas. Among the direct, deregulated target genes are classic HES1 target genes from other organ systems such as ATOH1 and *ASCL1*. Together, our data suggest a multi-layered mechanism for HES1 suppression of pancreatic endocrine development and additionally serves to suppress inappropriate expression of a set of non-pancreatic bHLH factors.

## Results

### Generation of *HES1* and *NEUROG3* mutant cell lines

To better understand the mechanism by which HES1 regulates pancreas development we took advantage of recent developments in directed differentiation of human embryonic stem cells (hESCs) to the pancreatic endocrine lineage via a series of progenitor stages (Figure 1 and (Rezania et al., 2014). Using the iCRISPR platform (González et al., 2014), we introduced indels in the HES1 and NEUROG3 genes, either singly or in combination in iCRISPR H1 cells.

**Figure 1:**
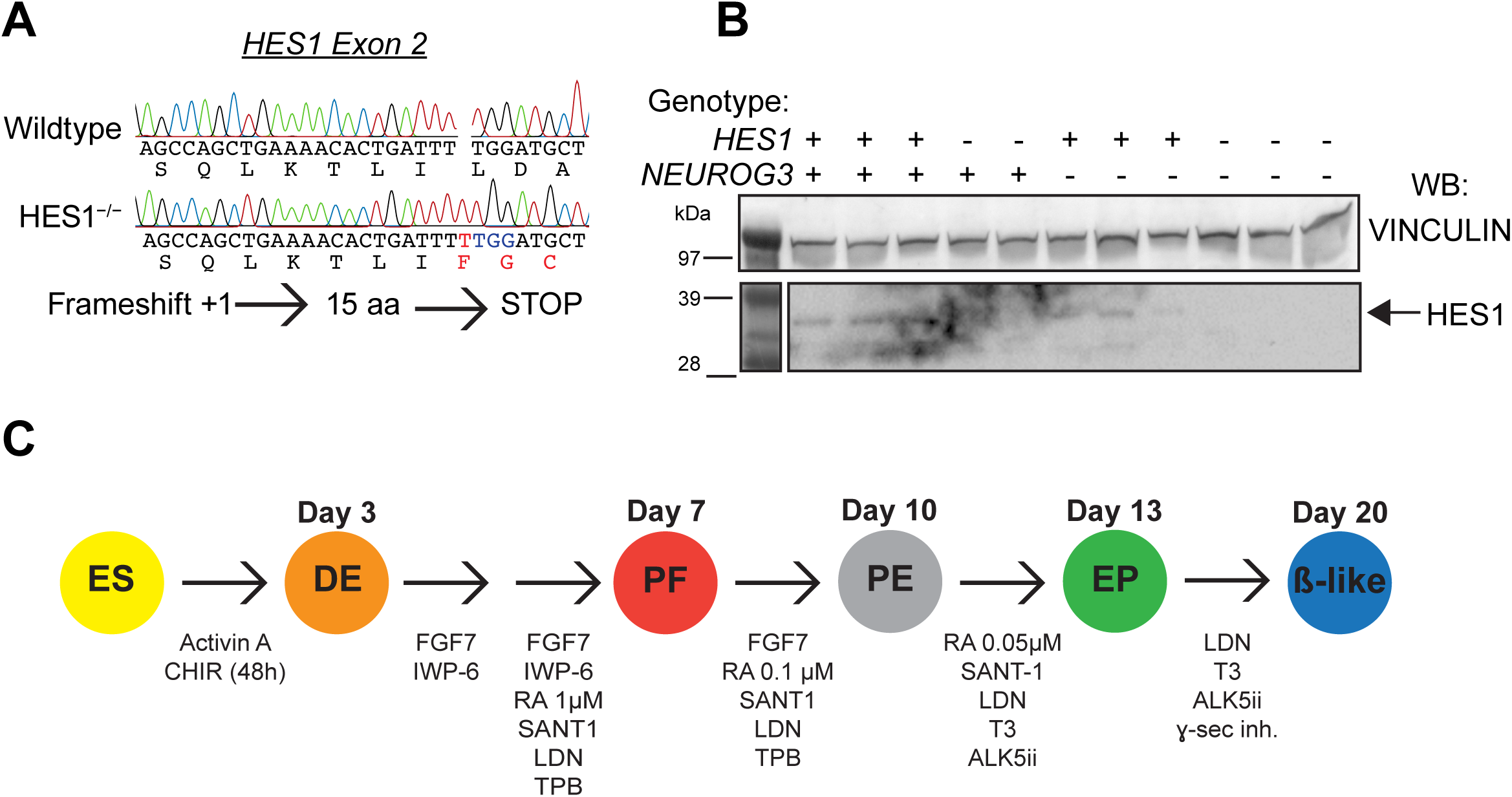
CRISPR-Cas9 mediated knock-out cell lines. A) Representative Sanger-sequencing trace from a PCR of the HES1 exon 2 around the cut site (PAM indicated in blue), showing introduction of a T indel (indicated in red). B) Western blot of clonal H1-iCRISPR lines. HES1 is not expressed in cell lines genotyped to be HES1^-/-^ (shown as -) or in HES1^-/-^NEUROG3^-/-^. C) Schematic of differentiation protocol to β-cell like cells.

Using previously described gRNAs (Zhu et al., 2016) to target exon2 of the NEUROG3 gene and exon2 of the HES1 gene we detected by Sanger and RNA-sequencing a GA insertion and a T insertion, respectively, resulting in premature STOP codons (Figure 1A and Sup. Figure 2). We confirmed the loss of HES1 protein by western blotting (Figure 1A,B) in two/multiple clonal cell lines carrying the introduced mutation and the loss of NEUROG3 by immunostaining (Figure 1B and Figure 2).

**Figure 2:**
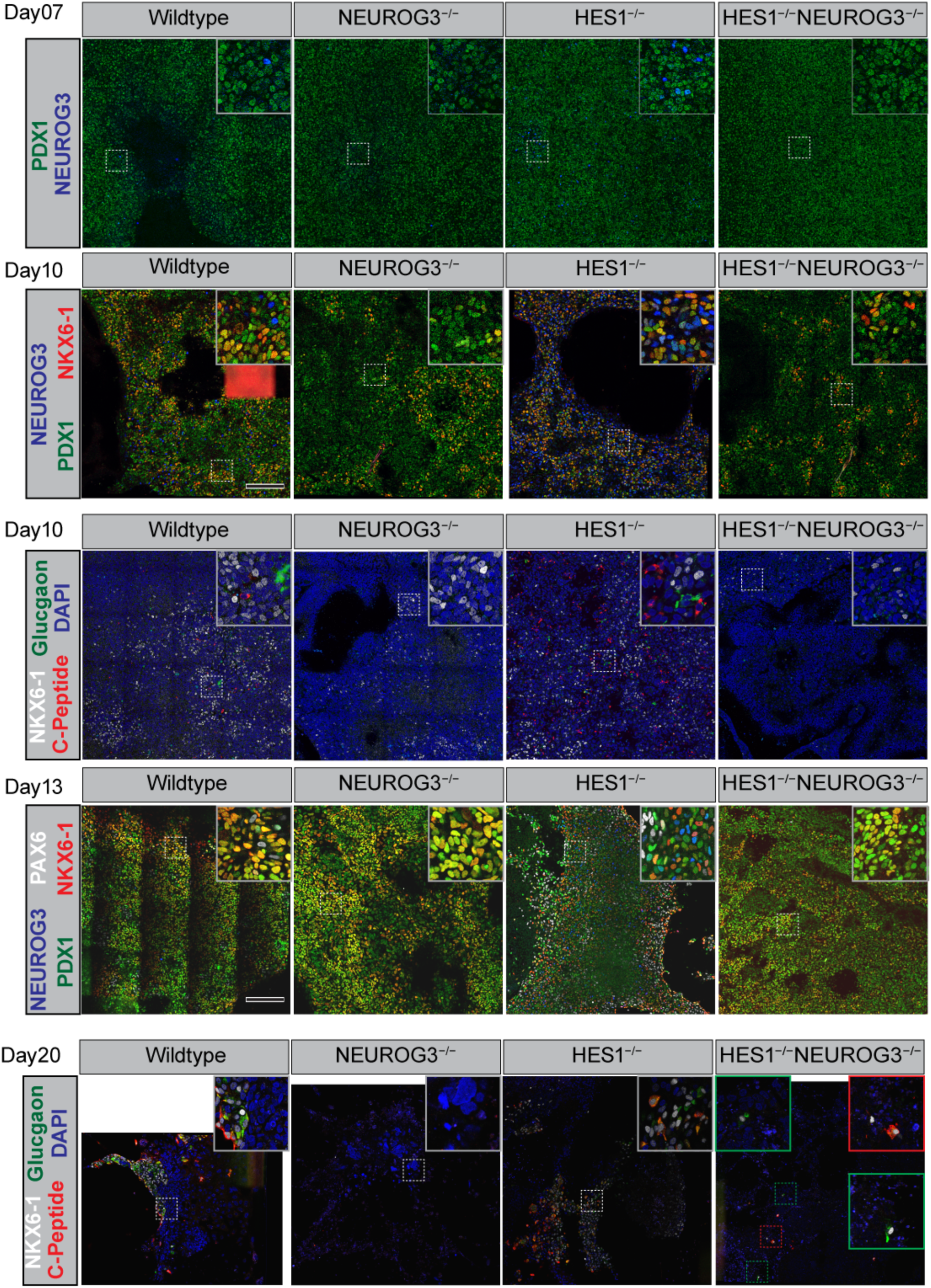
Immunostaining of hES cells differentiation towards pancreatic endocrine lineage for 10, 13 and 20 days. Black size bars indicate 200µm in the overview and same bar indicate 55µm in the insert.

We then subjected two or more clonal lines of H1-iCRISPR (wildtype), *HES1*^-/-^, *NEUROG3*^-/-^ and *HES1*^-/-^*NEUROG3*^-/-^ (abbreviated *H1N3*-dKO or *H1*^-/-^N3^-/-^) to differentiation to β-like cells using a modified version of the protocol from Rezania et al. (2014). Marker analysis showed that all cell - lines maintained pluripotency (*OCT4*^+^) as hESC and were able to differentiate to *FOXA2*^+^) and SOX17+ co-positive definitive endoderm (DE, Day 3), *PDX1*^+^) cells at the posterior foregut stage (PF, Day 7), to bipotent progenitors marked by *PDX1*^+^) and *NKX6-1*^+^) at the Pancreatic Endoderm stage (PE, Day 10), and the Endocrine Precursor stage (EP, Day 13) (Figure 2, Sup. Figure 8 and Sup. Figure 7).

### Precocious differentiation of the endocrine lineage in *HES1*^-/-^ cell lines

In a previous report, one of us showed an excess of C-peptide expressing cells at the polyhormonal-β stage in differentiated *HES1*^-/-^ cell lines (Zhu et al., 2016) consistent with a repressive role for HES1 on endocrine differentiation (Jensen et al., 2000b). Assessing the timing of endocrine commitment in more detail allowed us to extend the previous observation and demonstrate precocious NEUROG3+ cells in the *HES1*^-/-^ cell lines as early as Day 7 at the PF stage, similar to the precocious appearance of these cells in the E8.5 mouse embryo (Ahnfelt-Rønne et al., 2012; Jensen et al., 2000b). At the Day 10 PE-and Day 13 EP-stages, we observed a prominent increase of NEUROG+ cells in the *HES1*^-/-^ cell lines compared to wildtype while they were absent from the *NEUROG3*^-/-^ and *HES1*^-/-^*NEUROG3*^-/-^ cell lines as expected (Figure 2 and Sup. Figure 7).

C-peptide and glucagon single-positive cells were observed already at Day 10 and more prominently at Day 13 in the *HES1*^-/-^ cells, demonstrating very early differentiation to monohormonal α-and β-like cells (Figure 2 and Sup. Figure 7). This is also apparent by RT-qPCR, where GCG, INS, and NEUROD1 expression was markedly increased in *HES1*^-/-^ samples at both Day 10 and 13 compared to wildtype and with undetectable levels in the *NEUROG3*^-/-^ cell lines (Sup. Figure 8). In contrast to some of our previous work by Zhu and co-workers who saw around 0.5% C-peptide+ cells in Day 20 PH-β stage *NEUROG3*^-/-^ HUES-8 cells (Zhu et al., 2016), in present study we did not detect C-peptide by immunocytochemistry in NEUROG3^-/-^ H1 cells at the PH-β stage, however in the *HES1*^-/-^*NEUROG3*^-/-^ H1 cells they were present at a low frequency (Figure 2).

### RNA-sequencing reveals NEUROG3-dependent and -independent changes in the *HES1*^-/-^ transcriptome

To study the genome-wide transcriptional changes occurring upon loss of HES1 we performed RNA-sequencing (RNA-seq) on two differentiation experiments with two cell lines of each genotype on Day 13. Principal Component Analysis (Figure 3A) revealed that the samples and experiments cluster according to genotype and that *HES1*^-/-^ differs most from the other genotypes at this stage. We did differential expression analysis at the gene level using DESeq2 and found 1198 genes are significantly differentially expressed and upregulated in HES1^-/-^ cells compared to wildtype controls with an adjusted p-value ≤0.1 and log2 fold change ≥ 1.5. Similarly, we find 482 genes upregulated in *HES1*^-/-^*NEUROG3*^-/-^ over *NEUROG3*^-/-^ (Figure 3C).

**Figure 3:**
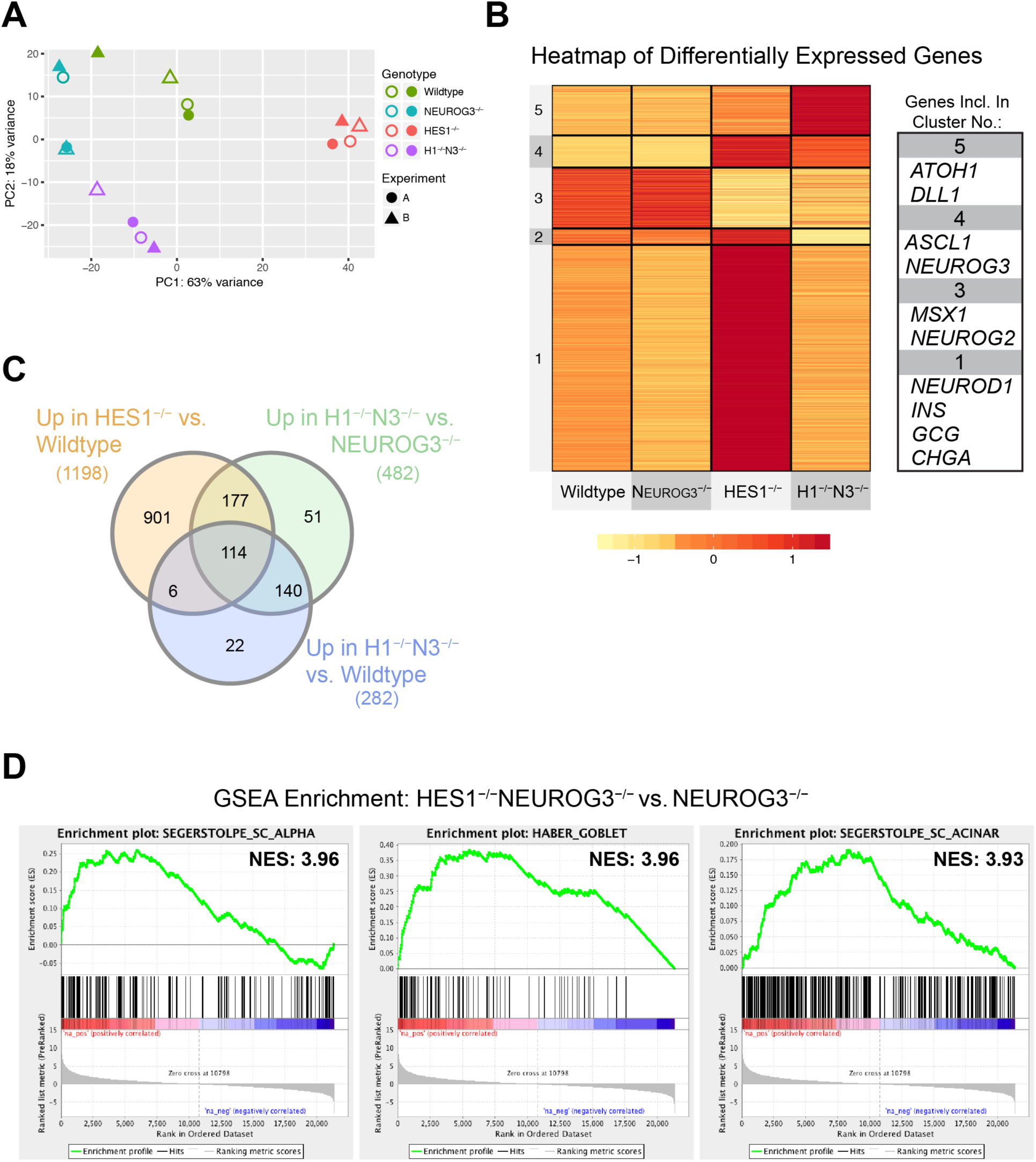
RNA-sequencing of Day 13. A) Principal Component Analysis, color-coding indicates genotype as shown in the legend, shape (triangle or round) indicate which experiment and filled or non-filled indicate the cell line. B) Heatmap of all differentially expressed genes by kmeans clustering analysis and scaled by row to show patterns of expression. Gene name of representative genes in the indicated clusters. C) Venn-diagram of the comparisons indicated by color-coding. D) GSEA enrichment of HES1^-/-^NEUROG3^-/-^ vs. NEUROG3^-/-^ with Normalised Enrichment Score (NES) for comparison. Black bars show gene position from the gene set in the ranked gene list (high in HES1^-/-^NEUROG3^-/-^ on the left). Green line shows the Enrichment Score based on overrepresentation of genes in the top of the ranked list. From the left, Alpha-cell signature from Segerstolpe et al., Goblet-cell from Haber et al. and Acinar-cell signature from Segerstolpe et al.

### Deregulation of endocrine lineage genes

To comprehend the differences in expression between the genotypes we did k-means clustering analysis on all genes that are deregulated in any comparison and this revealed succinct expression patterns for each genotype. Genes that are highly expressed in HES1^-/-^ compared to the other genotypes are clustered in group 1 and 2, and as expected they include peptide hormone genes such as *INS, GLU, SST, GHRL*, and *PYY* as well as numerous genes encoding endocrine-specific transcription factors such as *NEUROG3, NEUROD1, NEUROD4, PAX4, PAX6, INSM1*, and *ISL1*. Next, we performed Gene Set Enrichment and found in the HES1^-/-^ vs wildtype the top three datasets to be a very strong enrichment of the gene signature from “glucagon producing α-cells” (Normalised Enrichment Score, NES 6.62), “Oxididative phosphorylation” from the KEGG database and a gene signature of “Enteroendocrine cell” (Sup. Figure 6). Emphasising that the HES1^-/-^ cells strongly differentiate towards the endocrine lineage.

Interestingly, in a parallel analysis in HES1^-/-^NEUROG3^-/-^ vs. NEUROG3^-/-^ also found the signature “Alpha-cells”, showing that a significant set of genes expressed in α-cells are upregulated in the HES1^-/-^NEUROG3^-/-^ despite the loss of NEUROG3 (Figure 3), suggesting HES1 regulates certain endocrine lineage genes independently of NEUROG3. Remarkably, the “Goblet cell” signature from the intestinal system was also highly enriched driven by a number of goblet cell markers such as GFI1, *TFF3*, *KLF4* and *ATOH1*.

### Deregulation of Notch pathway genes

Notably, we find several genes encoding Notch pathway components among the genes that are deregulated by loss of HES1. Most significantly, the ligand-encoding genes *DLL1* and *DLL4*, as well as *NEURL1*, encoding an E3 ubiquitin ligase with several Notch ligands among its substrates (Koutelou et al., 2008) and MFNG, which encodes a glycosyltransferase that extends O-fucose moieties on Notch receptors (Kakuda and Haltiwanger, 2017), were upregulated in *HES1*^-/-^ cells compared to wild type, as well as in in *HES1*^-/-^*NEUROG3*^-/-^ cells compared to *NEUROG3*^-/-^ cells (**Error! Reference source not found**.). Genes showing more modest changes in expression upon loss of HES1, either in wildtype or on the NEUROG3^-/-^ background, include DLL3 (log2FC=1.39/1.68; p=0.0007/0.0001), *NOTCH1* (NC/1.24; NA/0.0006), *NOTCH2* (log2FC=−0.63/NC; p=4.32E-05/NA), *NOTCH3* (log2FC=−1.15/NC; p=6.97E-09/NA), *HEY1* (log2FC=1.11/0.76; p=0.00057/0.068), HEYL (log2FC=NC/1.14; p=NA/0.051), HES5 (log2FC=NC/2.54; p=NA/0.0006), *HES6* (log2FC=1.34/1.51; p=8.24E-06/1.81E-06), and LFNG (log2FC=1.52; p=0.024), which encodes another glycosyltransferase extending O-fucose moieties on Notch.

Remarkably, among the deregulated genes that are known targets of the Notch pathway we find not only genes expected to be expressed in the pancreas but also some that are not normally expressed in the pancreas. For example, we find significantly higher levels of *ASCL1*, *ASCL2* and *ATOH1*, transcripts that encode bHLH factors that are not expected to be expressed in the pancreas epithelium, in *HES1*^-/-^*NEUROG3*^-/-^compared to wild type or NEUROG3^-/-^ cell lines and confirmed for *ASCL1* by RT-qPCR (Sup. Figure 8). Interestingly, *HES1* mRNA remains unchanged in all genotypes despite the frame-shift mutation which would often lead to non-sense mediated decay, however, the potential loss of repression by HES1 protein on its own promoter could compensate by increasing gene expression (Takebayashi et al., 1994) (Kobayashi et al., 2009b; Sasai et al., 1992). *NEUROG3* mRNA levels are noticeably lower in the *NEUROG3*^-/-^ lines, but wild type levels are restored in the *HES1*^-/-^*NEUROG3*^-/-^ cell lines (Sup. Figure 8).

### HES1 does not regulate cell cycle genes

HES1 has been associated with regulation of the cell cycle through direct repression of the CDK inhibitor p57^KIP2^ (encoded by *CDKN1C*) (Georgia et al., 2006). However, we do not see HES1 binding to this gene, nor do find *CDKN1C* to be significantly deregulated between NEUROG3^-/-^ and HES1^-/-^*NEUROG3*^-/-^, suggesting that it is not a direct target gene (Sup. Figure 3). Only in the HES1^-/-^ samples is it upregulated compared to wild type pointing towards an indirect regulation of the gene, in that many cells enter a post-mitotic endocrine cell fate after loss of HES1. Also, we examined other well-studied proliferation markers such as *PCNA* and *MKI67* and found them to be downregulated only in the HES^-/-^ samples (Sup. Figure 3C). Assessing all cell cycle associated genes (GO term Cell Cycle) we found many to be deregulated after loss of HES1, but not in the *HES1*^-/-^*NEUROG3*^-/-^ cells. The few genes that are actually deregulated and in the GO term Cell Cycle are to our knowledge are not controlling cell cycle but are expressed in cell types that are generally not cycling, such as *NEUROD1* and *INSM1* in endocrine cells. Others such as *CDK5R2* is an activator of the atypical cyclin kinase *CDK5* and not regulating normal cell cycle (Martin et al., 2012).

### ChIP-seq identifies direct HES1 target genes

To identify genome-wide, direct targets of HES1 in human pancreas progenitors, we performed Chromatin immunoprecipitation followed by deep sequencing (ChIP-seq) of endogenous HES1 in the HUES4 *PDX1*^FP/+^ reporter cell line on Day 13 of differentiation from both unsorted bulk populations and FACS-sorted GFP+ cells. Peak finding by MACS2 revealed 998 peaks shared between the two replicates of unsorted Day 13 cells, distributed with 744 peaks near promoters, defined by 2kb upstream or 0.5kb downstream of transcription start sites (TSS), and 254 at distal sites (non-TSS). Motif search on 200bp around the summit of the 744 peaks close to TSS revealed an extended motif similar to the previously described HES/Hairy binding sites, the N-Box and the Class C site (Ohsako et al., 1994): a **SGCRCGYGC** motif which was contained in 64% of the peaks, indicated with red marks in Figure 4. Motifs are often found close to the peak centre such as for *DLL1* locus, but there are also matching motifs in the near vicinity of peaks, apparently not bound by HES1, suggesting other co-factors and the epigenetic environment determines the binding.

**Figure 4:**
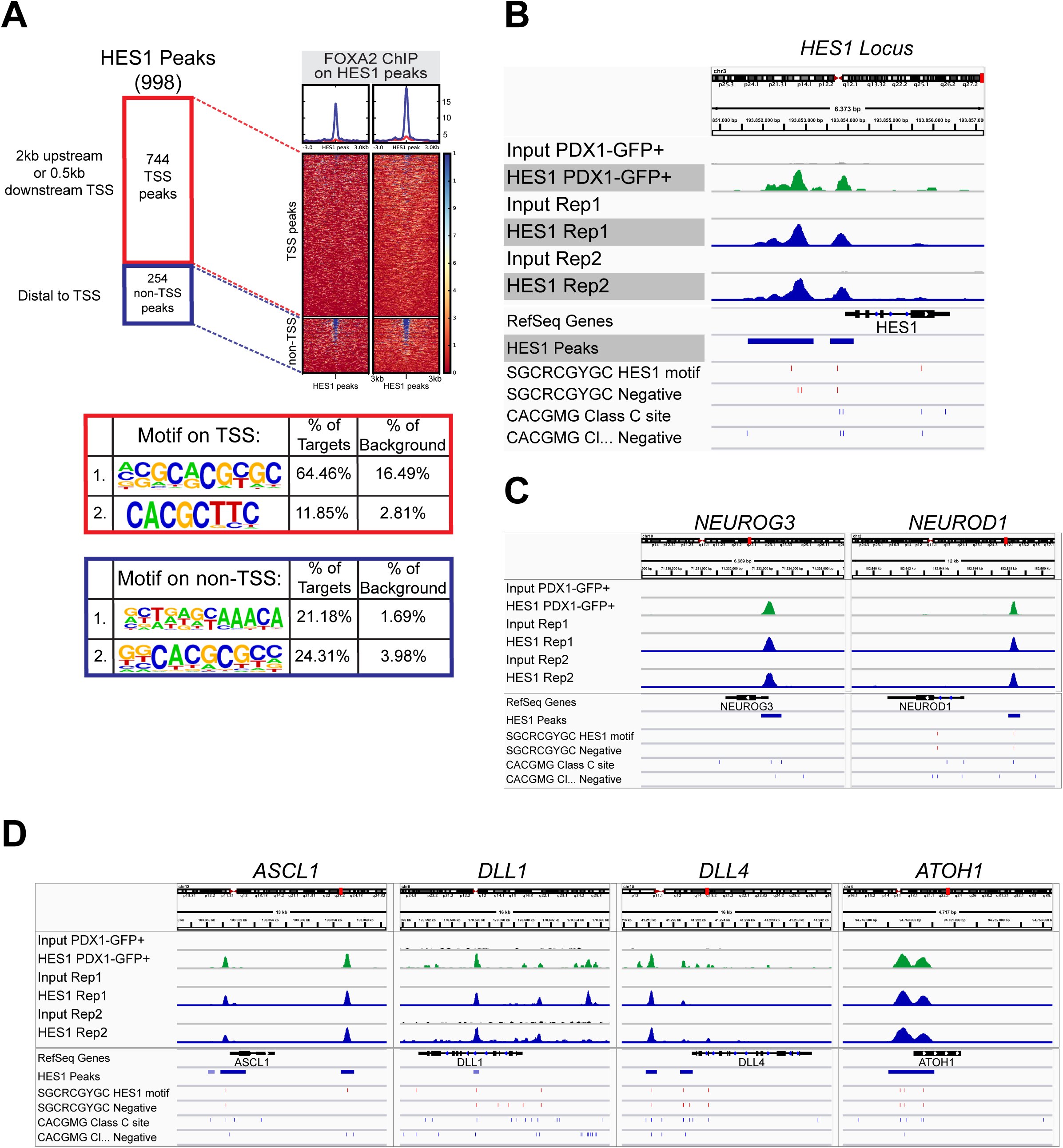
ChIP-seq of HES1. A) Distribution of peaks on TSS (within 2kb up and 0.5kb downstream of a TSS). Waveplot of FOXA2 binding to HES1 peaks. Homer de novo motif search shown in tables. B) ChIP-seq enrichment at HES1 locus. HES1 peaks row indicate peaks found by MACS2 in blue and below HES1 motif on each strand and Class C sites on each strand C) NEUROG3 and NEUROD1 loci D) ASCL1, DLL1, DLL4 and ATOH1 loci.

*NEUROG3* is an example of a bound gene which does not contain the full HES1 motif but does contain the core motif of the class C site: *cgcCACGCGag.*

Surprisingly, at the distal sites (non-TSS) we found a motif belonging to FOXA-family of transcription factors to be the most enriched, not the expected HES1 motif. HES1 has to our knowledge not previously been associated with FOXA2 and we took advantage of our previously published FOXA2 ChIP-seq data (Weedon et al., 2014) to show an abundance of FOXA2 localised to a subset of the non-TSS peaks but not to the peaks located close to TSS.

Identification of HES1 bound genes that are also deregulated in *HES1*^-/-^*NEUROG3*^-/-^ compared to *NEUROG3*^-/-^ samples yielded 28 genes that are de-repressed (Figure 5). The mean expression across genotypes are shown as a heat map in Figure 5C, which reveals several genes expressed in the endocrine lineage and previously described to be downstream of NEUROG3, including *DLL1*, *DLL4*, *MFNG*, *NEUROD1*, *MNX1*, *INSM1* and *NEUROG3* itself (Gradwohl et al., 2000) (Schwitzgebel et al., 2000) (Jensen et al., 2000a) (Gasa et al., 2004), which are all upregulated in D13 EP stage *HES1*^-/-^*NEUROG3*^-/-^cells in spite of the absence of NEUROG3. Although DLL1 is likely not to be endocrine-specific (Ahnfelt-Rønne 2012), these data nevertheless show that loss of HES1 is sufficient to promote the expression of a number of genes expressed specifically in the endocrine lineage independently of the presence of NEUROG3 protein. This unexpected finding is revealing that HES1 represses endocrine lineage determination at multiple levels.

**Figure 5:**
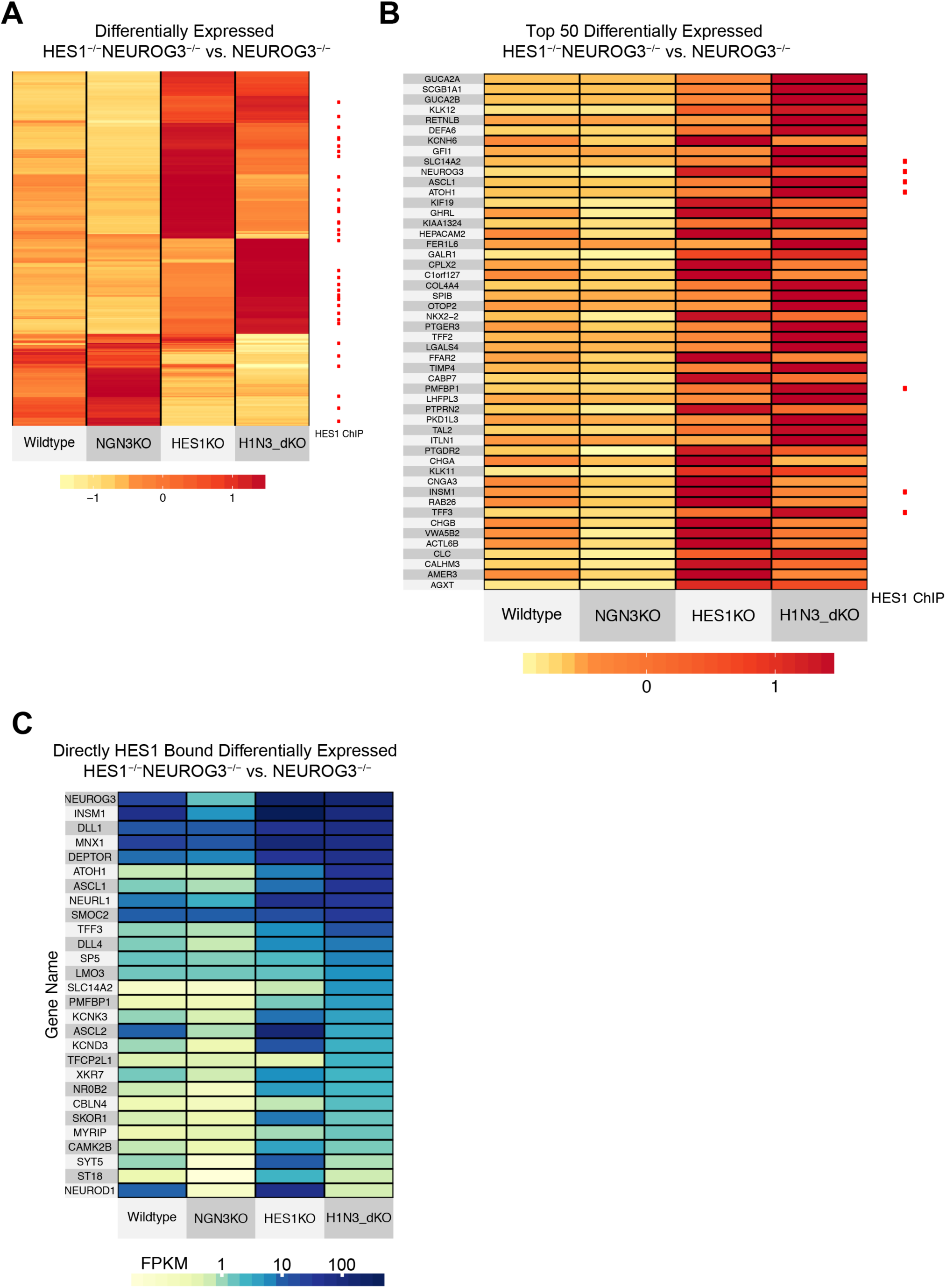
Differentially expressed genes upon HES1 loss that are also directly bound by HES1. A) A subset of differentially expressed genes between HES1^-/-^NEUROG3^-/-^ and NEUROG3^-/-^ are bound by HES1 ChIP (red dots). B) Top 50 upregulated genes shown in the heatmap with HES1 bound genes indicated with red dots. C) 28 genes are bound by HES1 and deregulated in HES1^-/-^NEUROG3^-/-^ and NEUROG3

A number of genes, including *PAX6*, *CCK*, and *DUSP4* are upregulated in *HES1*^-/-^ cells compared to wildtype, but not in *HES1*^-/-^*NEUROG3*^-/-^ cells compared to *NEUROG3*^-/-^ samples, suggesting that activation of these genes depends strictly on NEUROG3. Given that they are bound by HES1 it is possible that they also depend on HES1 being downregulated, but that it is not sufficient. Unfortunately, our current experimental setup does not allow us to test for this.

Several genes containing FOXA2 and HES1 bound sites are not normally expressed in pancreatic tissue. These include homeobox genes such as *CDX2* and *ISX* as well as bHLH genes such as *ASCL1*, which are expressed in other endodermal organs like the intestine or, in the case of *ASCL1*, the lungs and stomach. Both ISX and *ASCL1* are upregulated in *HES1*^-/-^ compared to wild type samples and also in *HES1*^-/-^*NEUROG3*^-/-^ compared to *NEUROG3*^-/-^ samples, suggesting that HES1 may prevent inappropriate expression of these genes in the pancreas (**Error! Reference source not found**.).

Lastly, we compared our HES1 ChIP-seq data to the previously published ChIP-on-CHIP of Hes1 in mouse embryonoic stem cells from the Kageyama lab (Kobayashi et al., 2009b) by liftover of the reported peak to the human genome (hg19) and found HES1 enrichment on half of the reported top 50 peaks (Sup. Figure 4). This observation shows a remarkable conservation of HES1/Hes1 binding across species and interestingly also cell types, highlighting the relevance of present HES1 binding studies for other systems.

## Discussion

Here we have uncovered novel functions of HES1 in pancreas development and unravelled a multi-layered mechanism that secures robust regulation of pancreatic endocrine development. Our work highlights the importance of understanding the transcription factor networks that are regulating human β-cell differentiation and for development of efficient and precise protocols that enable the generation of β-like-cells suitable for diabetes therapy. Taking advantage of recent advances in protocols for deriving β-like-cells from hESCs (Pagliuca et al., 2014; Rezania et al., 2014) we used the iCRISPR platform to induce mutations in the *HES1* and *NEUROG3* genes in order to understand the molecular mechanisms of HES1-regulated pancreatic cell fate decisions.

Consistent with a previous report on the HUES4 cell lines (Zhu et al., 2016), the H1-iCRISPR *HES1*^-/-^ cell lines exhibit excessive endocrine differentiation, and in line with previous publications in mice, we find precocious appearance of the early endocrine progenitor marker NEUROG3 at Day 7 and of c-peptide and glucagon as early as Day 10 on both protein and mRNA level (Ahnfelt-Rønne et al., 2012; Jensen et al., 2000b). Beyond single markers we used GSEA to assess enrichment of gene signatures in *HES1*^-/-^ cells and as expected found genes expressed in α-cells as defined by single cell RNA-sequencing experiments (Segerstolpe et al., 2016) as well as Oxidative Phosphorylation from the KEGG database.

While expected, the huge shift in cell fate, towards endocrine differentiation, that is provoked by HES1 deficiency prevented the identification of genes that potentially were deregulated specifically in the progenitor population, where HES1 is active. We therefore used a “biological filter” to avoid the massive secondary transcriptional changes associated with the endocrine lineage choice by taking advantage of the impaired endocrine differentiation of NEUROG3 null cells and compared *HES1*^-/-^*NEUROG3*^-/-^ double knockout-to *NEUROG3*^-/-^ single knockout cells, which allowed us assess NEUROG3 independent HES1 target genes. A similar approach has previously been used in the Shivdasani lab to “filter out” specific intestinal cell populations (Kim et al., 2014).

We analysed potential changes in cell cycle regulation. Hes1 has previously been proposed to regulate the cell cycle in embryonic mouse pancreas through binding to an E-box in the Cdkn1c gene and thereby directly repress p57^Kip2^ expression (Georgia et al., 2006), and others have suggested that sustained expression of Hes1 inhibits cell cycle progression through repression of *PCNA*, *E2F1*, and *CDKN1A* (encoding p21^CIP1^) (Baek et al., 2006; Castella et al., 2000; Georgia et al., 2006; Hartman et al., 2004; Hatakeyama, 2004; Murata et al., 2005; Ström et al., 2000). However, although we did observe increased *CDKN1C* expression in the *HES1*^-/-^ cells compared to wildtype, we observed no change when comparing *HES1*^-/-^*NEUROG3*^-/-^ to *NEUROG3*^-/-^ cells, suggesting indirect activation of *CDKN1C* in endocrine committed cells downstream of NEUROG3 and not a direct effect of HES1 deficiency. Consistent with this notion, we did not observe HES1 binding to the *CDKN1C* gene in our ChIP-seq data. Similarly, the decrease in *PCNA* and *MKI67* expression argues against HES1-mediated repression of these genes. Indeed, *PCNA* and *MKI67* expression was restored in *HES1*^-/-^*NEUROG3*^-/-^ cells suggesting that this pattern can be explained simply by the increased proportion of post-mitotic NEUROG3^+^ and PAX6^+^ cells, and the associated depletion of proliferating PPs seen in our Day 13 *HES1*^-/-^ hESC-PP cultures.

From our HES1 ChIP-seq data, which is to our knowledge the first high quality ChIP-seq analysis of endogenous HES1, we identified 998 high confidence peaks. De novo motif search yielded the sequence “SG**CRCGYG**C”, which contains the Class C Hairy binding site in the majority of the peaks located close to a TSS (Ohsako et al., 1994; Sasai et al., 1992; Takebayashi et al., 1994).

Unexpectedly, we found a FOXA motif as the most enriched on distal regulatory elements, and an analysis of FOXA2 ChIP-seq data from hESC-PPs previously published by one of us (Weedon et al., 2014), revealed FOXA2 binding close to many of these HES1 peaks. It is possible that FOXA2, a known pioneer factor (Iwafuchi-Doi et al., 2016), acts to open the chromatin allowing HES1 to access its binding sites. It should be stressed however that we cannot be certain that co-binding occurs in individual cells. Nevertheless, many of the genes bound by both HES1 and FOXA2 are part of complex transcription factor networks that regulate cell fate in other endodermal tissues. This includes *CDX2*, *ISX* in the intestines and *ASCL1* in the stomach and lungs (Imayoshi et al., 2013; Ito et al., 2000; Jensen et al., 2000b; Kobayashi et al., 2009b; Kokubu et al., 2008) and it is likely that expression of such genes would interfere with normal pancreas development. In our hESC-PP cultures it is noteworthy that intestinal markers such as *IHH* and *VIL1* are induced, albeit modestly, by loss of *HES1*, in both wildtype and *NEUROG3* deficient background, suggesting a shift towards intestinal fate (**Error**! **Reference source not found**.).

Previously, the Kageyama lab published a ChIP-on-chip analysis of Hes1 binding sites in mouse embryonic stem cells (Kobayashi et al., 2009b). By comparison we found half of the reported peaks (top 50) to be enriched for HES1 on the corresponding site by doing lift-over from mouse genome (mm7) to human genome (hg19) (Sup. Figure 4). This shows conserved binding across species and strikingly across cell types suggesting HES1 binds many of the same genes independent of cellular context but rather that additional repressors and activators decide the contextual output. Among the Hes1/HES1 target genes shared between hESC-PPs and mESCs is *Hes1/HES1* itself. As shown in Figure 4, we observe HES1 binding to several sites in its own promoter suggesting that, similar to many other tissues, HES1 may regulate its own promoter activity in a negative feedback loop in pancreas progenitors. This negative feedback is key to ultradian oscillations in Hes1 promoter activity and Hes1 protein expression observed in mouse embryos in presomitic mesoderm as well as in neural progenitors (Hirata et al., 2002; Shimojo et al., 2008). We are currently investigating whether ultradian oscillations are a feature of pancreatic Hes1 expression and function.

Knock-out of HES1 on a NEUROG3-null background yielded 482 de-repressed genes (>1.5 log2FoldChange) of which 28 genes contained identified HES1 ChIP-seq peaks, these genes included an array of endocrine markers such as *NEUROD1* and *INSM1*, two key transcription factors previously thought to be downstream of NEUROG3. *PAX6*, encoding another well described endocrine transcription factor, is also bound by HES1, but its expression appears to depend strictly on NEUROG3. It his appears that HES1 regulates endocrine differentiation by repressing the key endocrine-inducing TF NEUROG3 as well as some of its target genes, a certain subset of which depends on HES1 activity in order not to become prematurely expressed. HES1 thus employs a multi-layered mechanism to regulate endocrine differentiation.

HES1 bound and regulated several Notch pathway genes, including the ligand encoding genes *DLL1* and *DLL4*. Except for a ChIP-on-CHIP experiment showing Dll1 and Dll4 to Hes1 bound by Kobayashi and co-workers (Kobayashi et al., 2009a), Dll1 has been described to be induced by bHLH factors such as Ascl1 and Neurog2/3 (Bel-Vialar et al., 2007; Castro et al., 2006; Lacomme et al., 2012) (Gasa et al., 2004; Lacomme et al., 2012; Treff et al., 2006), but we show here that DLL1 and DLL4 are not only bound by HES1, but also derepressed when *HES1* is deleted. Crucially, derepression was also observed in the absence of NEUROG3 suggesting that other transcription factors are responsible for their activation. Consistent with this notion, *DLL1* and *DLL4* were recently found to be bound by SOX9 and PDX1 in hESC-PPs (Shih et al., 2015; Wang et al., 2018). We also find the E3 ubiquitin ligase *NEURL1* to be upregulated in the HES1^-/-^NEUROG3^-/-^. *NEURL1* is an ortholog of Drosophila *Neuralized* which is necessary for Notch ligand activation in flies, but its function in mammals is not well understood as, in contrast to the fly, Neurl1^-/-^ and Neurl1/2 double knockout mice are viable and fertile with only minor phenotypes (Koo et al., 2007; Ruan et al., 2001; Vollrath et al., 2001). Further investigations are needed to determine if NEUROG3 directly regulates Notch ligands in the pancreas or if lateral inhibition is determined by HES1-Delta/Jagged-Notch feedback loops only.

Importantly, we also show HES1 binding to the *NEUROG3* promoter and that loss of HES1 is sufficient to activate NEUROG3 expression in hESC-PPs even without the auto-activating effect of NEUROG3 previously proposed to drive cells from a NEUROG3^Lo^ to a NEUROG3^Hi^ state in mice and *in vitro* (Ejarque et al., 2013; Gasa et al., 2004; Wang et al., 2008). Some of us has previously shown that some NEUROG3-independent endocrine cells are formed (~0.5% C-peptide^+^) by FACS analysis in the HUES4 iCRISPR NEUROG3^-/-^ cell line (Zhu et al., 2016), However, in the present study on the H1 hESC background we do not observe this by immunostaining, in line with other studies on NEUROG^-/-^ cells assessed by immunostaining (McGrath et al., 2015). Neurog3 knock-out mice were first described to be devoid of any endocrine cells but additional knock-out alleles on different mouse backgrounds have revealed small numbers of endocrine cells despite the loss of Neurog3 suggesting additional mechanisms to turn on the endocrine programme (Gradwohl et al., 2000; Wang et al., 2008). Also, occasional glucagon+ cells are observed in *Dll1*^-/-^*Neurog3*^-/-^ and *Hes1*^-/-^*Neurog3*^-/-^ mouse embryos (Ahnfelt-Rønne et al., 2012; Jørgensen et al.). Consistent with such mouse data, we observe some NEUROG3-independent endocrine cells in the *HES1*^-/-^*NEUROG3*^-/-^, indicating that HES1 could additionally de-repress other factors capable of driving endocrinogenesis although at a very low frequency. In zebrafish the *ASCL1*-ortholog ascl1b, rather than neurog3 control pancreatic endocrine cell fate, indicating that other bHLH factors such as ASCL1 or ATOH1 which are highly upregulated in the *HES1*^-/-^*NEUROG3*^-/-^ could potentially drive endocrine cell fate choice, as they do for stomach and intestinal endocrine cells, respectively (Flasse et al., 2013; Kokubu et al., 2008; Yang et al., 2001). Similar observations in vivo and lineage tracing experiments would be necessary to prove such plasticity of bHLH factors in the pancreas which was previously described in a neuronal system (Parras et al., 2002). Remarkably, *ASCL1* and *ATOH1* expression was much higher in *HES1*^-/-^*NEUROG3*^-/-^ cells than in *HES1*^-/-^ cells, showing that NEUROG3 is repressing expression of these two genes. B-class bHLH factors have previously been implicated in repressing each other’s expression. For example, Atoh1 and Neurog1 mutually inhibit expression of the other factor in the brain (Gowan et al., 2001) and Ascl2 blocks the myogenic activity of MyoD in muscle satellite cells by occupying MyoD binding sites in a non-productive fashion (Johnson et al., 1992) (Wang et al., 2017).

We find a “Goblet cell” signature enriched in the *HES1*^-/-^*NEUROG3*^-/-^ expression dataset by GSEA analysis including the HES1 bound gene *ATOH1* which is required for goblet cell formation in the intestine (Yang et al., 2001), which is in line with Notch loss of function mouse models Hes1^-/-^, Rbpj^-/-^and Dll1^-/-^Dll4^-/-^where Atoh1 is upregulated in the intestine leading to an increase in goblet cells (Jensen et al., 2000b; Pellegrinet et al., 2011). Furthermore, Notch overexpression in the intestine, and thereby sustained Hes1 leads to a failure to form goblet cells ((Fre et al., 2005; van Es et al., 2005)). Remarkably, expression of goblet cell markers is further increased in *HES1*^-/-^*NEUROG3*^-/-^ cells compared to *HES1*^-/-^ cells, and it is noteworthy that ectopic goblet cell formation is observed in the stomach of Neurog3^-/-^ mice. However, there has been no reported increase in goblet cell numbers in neither *Neurog3*^-/-^ nor *Hes1*^-/-^ pancreas. Although they receive little attention, mouse pancreatic ducts do contain mucin-producing goblet cells in addition to the classical bicarbonate-secreting principal cells (Lev and Spicer, 1965). These goblet cells are likely differentiating from the trunk progenitors, which then should be considered tri-potent rather than bi-potent. We hypothesise that upon low Notch activity a tri-potent progenitor would allow Atoh1 or Neurog3 upregulation leading to a goblet or endocrine lineage cell fate decision, respectively, possibly dependent on anatomical location, developmental stage, or the relative levels of Neurog3 and Atoh1. Alternatively, the goblet cell signature could be due to inappropriate activation of an intestinal transcriptional programme in *HES1*^-/-^ hESC-PP cells and particularly in *HES1*^-/-^*NEUROG3*^-/-^ hESC-PP cells and would shed light on the mechanisms by which Notch-initiated transcriptional networks and bHLH factors interplay in the endoderm.

Together, our results uncovered several new features of HES1-regulated pancreatic cell fate choice in developing hESC cultures. This knowledge can be used to refine differentiation protocols, but understanding how these findings translate to the embryonic pancreas in vivo requires further analysis of conditional Hes1 and Hes1/Neurog3 knockout mice.

## Materials and Methods

### Generation of knock-out human embryonic stem cell lines

H1 human embryonic stem cell lines with a doxycycline inducible Hygro-iCRISPR-Cas9 (iCRISPR) Wildtype (clone E8) and H1 Hygro-iCRISPR NEUROG3^-/-^ (clone N6_H3) were generated by Z. Zhu and D. Huangfu, Columbio University, similarly to previously described in HUES8 human embryonic stem cell line (González et al., 2014; Zhu et al., 2016).

Cells were maintained in the E8 media on human recombinant Vitronectin (ThermoFisher) according to manufacturer’s instructions changing media every day and splitting 1:10 using Versene (ThermoFisher) to dissociate for 5-7min, removing the Versene and lifting cells with fresh media by pipetting.

In vitro transcribed gRNA “Cr4” targeting HES1 exon2 from Zhu et al., by PCR amplifying template DNA oligo (IDT) to double stranded DNA with Phusion High Fidelity PCR kit (NEB) in a 50 µL reaction: 28.5 µL H2O, 5µL 10µM primer mix (F: *TAATACGACTCACTATAGGG*, R: *AAAAGCACCGACTCGGTGCC*), 10µL Phusion Buffer, 1µL 10mM dNTP, 0.5 µL Phusion enzyme and 5µL ssDNA oligo Cr4:

*TAATACGACTCACTATAGGG****CCAGCTGAAAACACTGATTT****GTTTTAGAGCTAGAAATAGCAAGTTA AAATAAGGCTAGTCCGTTATCAACTTGAAAAAGTGGCACCGAGTCGGTGCTTTT.*

Gel purified PCR-product was diluted to 100nM for T7 IVT reaction using MEGAshortScript T7 Transcription kit (ThermoFisher): 2 µL reaction buffer, 8µL dNTPs, 1µL dsDNA template, 2µL enzyme, 7µL H_2_O. After 4h incubation at 25°C, added DNAse 15min 37°C and purified with RNeasy Micro kit (Qiagen) to a concentration about 1µg/mL stored at −80°C. To evaluate transfection efficiency by microscopy, we also made a gRNA with Cy3-flourescence labeled dCTP, however the cells transfected with these was not used for further work.

Wildtype and NEUROG3 H1 hygro-iCRISPR cell lines were treated with doxycycline (1µg/mL) for 48 hours prior to transfection. Cells were washed twice with PBS and dissociated with TrypLE Select (ThermoFisher) for 5 minutes, diluted in media and spun down 4 min 200g. Resuspended in 1mL of media. Seeded 800.000 cells/well 1.5mL media in a 6-well. Made 150µL OptiMEM (ThermoFisher) with 9µL RNAiMAX (ThermoFisher) and mixed with 150µL OptiMEM with 1µg gRNA, incubated 5min and dropped on the cells. Changed every day and split cells three days after transfection to clonal density, around 2000 cells per 10cm plate. Used approximately 1/5 of the leftover cells for transfection efficiency genotyping: pelleted cells, removed media, added 20µL QuickExtract (EpiCentre) and incubated >3h at 65°C, then stopped the reaction at 95°C 15min. and diluted 1:10 for PCR reaction. PCR reaction was setup using Phusion HighFidelity MasterMix (NEB) 5µL, 1µL 10µM mix of primers Fwd: *CGGATAAACCAAAGACAGCA* and Rev: *CATAGAGTAGGCAAGAAAGGA*, 3% final volume DMSO, 2.7µL H2O and 1µL DNA. 98°C 4 minutes, 34 cycles of 98°C 30 sec., 60.8°C 30 sec., 2 min 72°C and then a final extension at 72°C 10 minutes. PCR reaction run on an agarose gel to check band size and purity and then gel purified with Centrifugal Unit for Gel Extraction (Millipore) and subsequently purified using DNA Clean and Concentrator-5 (Zymo) and eluted in 8µL. 5µL eluted PCR product with 5µL 5µM sequencing primer(*TGTATCTCTTTGCAGCCCCT*) was sent for Sanger Sequencing at GATC Biotech. CRISPR-Cas9 efficiency was estimated using TIDE software (https://tide.deskgen.com) (Brinkman et al., 2014) to be up to 65% indel formation.

Picked the number of clones accordingly, 10 days after clonal seeding in PBS to an 1.5mL tube containing 50µL versene for dissocation. Took approximately half for seeding in a 24-well and the other half for clonal PCR by same protocol as for the transfection efficiency PCR. Using TIDE software to estimate the genotype, skipping everything that potentially could be heterozygous and Topo-cloned the PCR product of the putative knock-out clones. Sanger-sequenced >6 Topo-Clones for each hES-clone and evaluated validity of the genetic knock-out also by western blot. HES1^-/-^

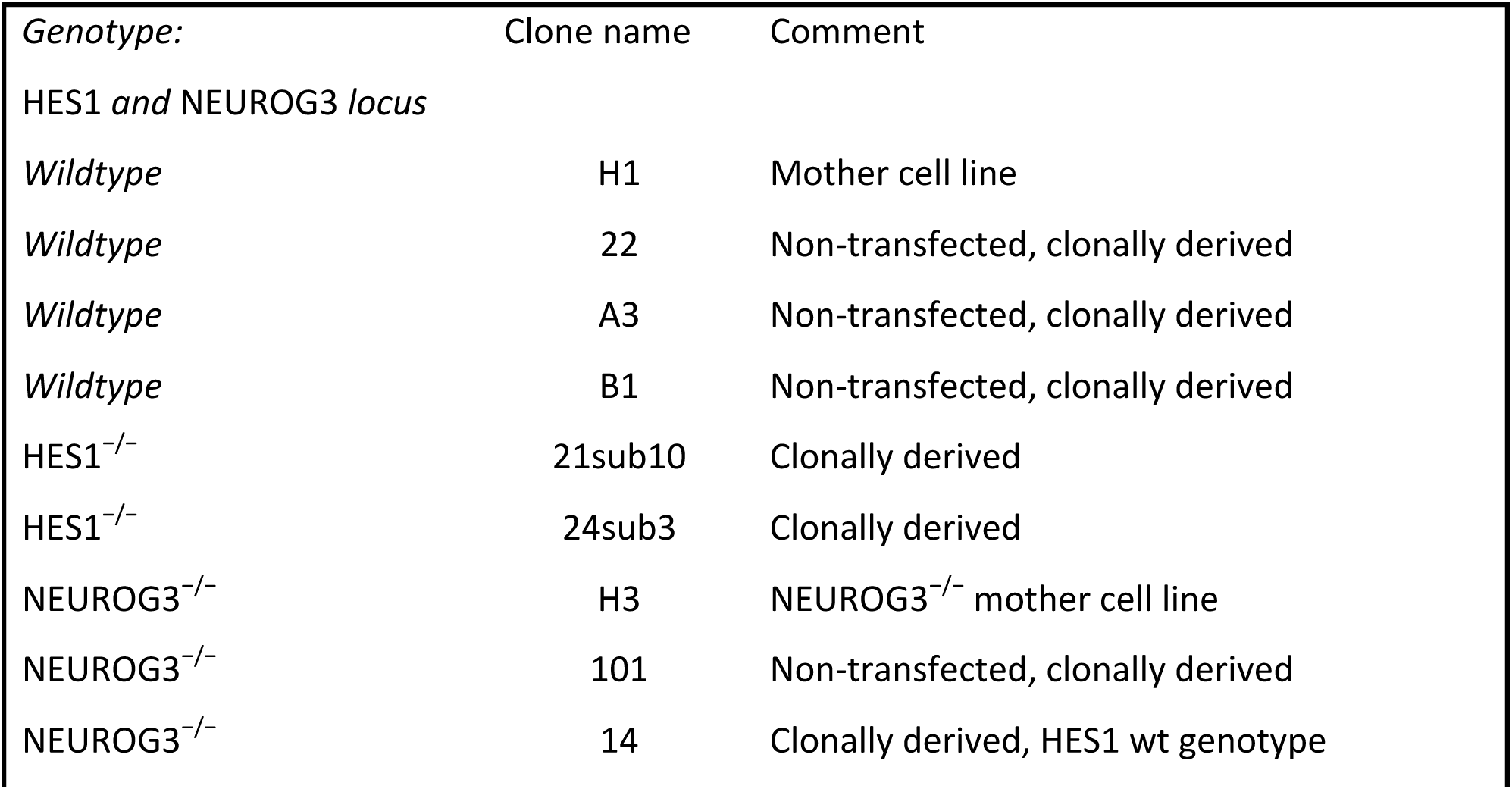
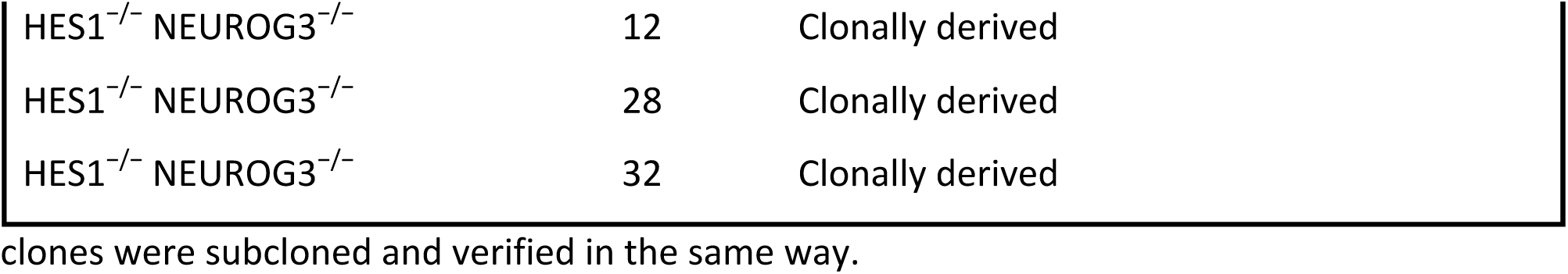

### Differentiation protocol of human embryonic stem cells to human pancreatic β-like cells

Seeded each cell line at 150.000cells/cm^2^ in 12-wells in 1mL and Ibidi µslides in 300µL after single cell suspension with TrypLE Select 5 min, spin down and resuspended in media then counting on Sceptor cell counter 40µM (Millipore) diluted 1:10 in PBS. Differentiation experiments were performed three times (Diff #3, #4 and #5), each for all 12 cell lines, harvested for RNA and protein on various days, including for RNA Day 10 and Day 13 for all experiments. For immunocytochemistry staining cell lines H1, 22, 21sub10, 24sub3, H3(4^th^ diff exp.), 101, 14(5^th^ diff. exp.), 12, 28 were seeded in Ibidi slides as explained above.

For 48 hours prior to differentiation cells were kept in E8 hESC media changing media every day also for differentiation. The protocol is slightly modified version from the Kieffer Lab(Rezania et al., 2014) including porcupine inhibitor IWP-6 at 5µM for days 3-6. Compounds and solutions are from ThermoFisher unless otherwise stated.

| Compound | Vendor | Cat # | Stock | unit | Reconstitution |
| --- | --- | --- | --- | --- | --- |
| BSA | Proliant | 68700 | 10 | % | PBS |
| CHIR 99021 | AxonMedChem | 1386 | 6 | mM | DMSO |
| Activin A | PreproTech | 120-14 | 100 | μg/mL | PBS |
| Ascorbic Acid | Sigma | A4544-25G | 250 | mM | 44mg/mL H2O |
| FGF7 (KGF) | Peprotech | 100-19 | 500 | μg/mL | 10ug in 20μL PBS (-80C) |
| Retinoic acid | Sigma | R2625 | 10 | mM | DMSO |
| Sant-1 | Sigma | S4572 | 2.5 | mM | 5mg in 5354μL DMSO |
| LDN | AxonMedchem | DM 3189 | 1 | mM | PBS |
| TPB (PKC act V) | EMD | 565740 | 1 | mM | 1mg in 1.95mL DMSO (-20) |
| Alk5i II | Sigma | R0158-5MG | 10 | mM | 5 mg in 1740 μL DMSO (-20°C) |
| T3 | Sigma | T6397 | 20 | μg/ml | 20μg/mL (29,72μM in MCDB-131) |
| Heparin | Sigma | H3149 | 50 | mg/mL | PBS |
| y-sec. Inh XX | EMD/ Merck | 565789 | 1 | mM | 500 μg in 1.079 ml DMSO |
| Y27632, Rock Inh | Sigma | Y27632 | 10 | mM | 1mg in 312 μL PBS: 10mM |
| Zinc sulphate | Sigma | Z0251 | 10 | mM | MCDB-131 |

S1-S2 Basal Media containing sodium bicarbonate, glucose, BSA and Glutamax in MCDB-131 media prepared in advance. In order to obtain Definite Endoderm cells at the end of Stage 1,compounds were added to make up Stage 1 (S1) media with the addition of CHIR and Activin A according to below tables for 2 days (t=0, t=24h), and for the 3^rd^ day not adding CHIR, but only Activin A(t=48h). For Stage 2 (S2) in the same basal media according to below scheme S2 Media to get Primitive Gut Tube (GT). See also scheme in (Figure 1)

| **S1-S2 basal media** |  |  |  |  |
| --- | --- | --- | --- | --- |
|  | Stock conc.: | stock unit | final conc.: | unit |
| Sodium bicarbonate |  |  | 1.5 | g/L |
| Glucose, 10mM, (1000mg/L already) | 200,000 | g/L | 1800 | mg/L Total |
| BSA, 0.5% | 0.1 | BSA | 0.5% |  |
| Glutamax | 100 | x | 1 | x |

| **S1 Media** for Definitive Endoderm (DE), t=0, t=24h, t=48h<br>(note Day2/48h only Activin A, no CHIR) |  |  |  |  |
| --- | --- | --- | --- | --- |
|  | Stock conc.: |  | final conc.: |  |
| CHIR, 3mM | 6,000 | μM | 3 | μM |
| Activin A, 100μg/mL | 100,000 | ng/mL | 100 | ng/mL |

| **S2 Media** Primitive Gut Tube (GT) **Day 3-4** |  |  |  |  |
| --- | --- | --- | --- | --- |
|  | Stock conc.: |  | final conc.: |  |
| Ascorbic Acid, 0.25mM | 250 | mM | 0.250 | mM |
| FGF-7, 500μg/mL stock | 500,000 | ng/mL | 50 | ng/mL |
| IWP-6, 5μM | 10,000 | μM | 5 | μM |

For Stages 3 and 4 (S3 and S4) medias were prepared to obtain Posterior Foregut (end of stage 3 / on Day 7) and Pancreatic Endoderm (EP) end of stage 4 on Day 10.

| **S3-S4 Basal media** |  |  |  |  |
| --- | --- | --- | --- | --- |
|  | Stock conc.: |  | final conc.: |  |
| Sodium bicarbonate |  |  | 2.5 | g/L |
| Glucose, 10mM, (1000mg/L already) | 200000 | g/L | 1800 | mg/L Total |
| BSA, 2% | 10% |  | 2.0% |  |
| Glutamax | 100 | x | 1 | x |

| Stage 3 Posterior Foregut (PF) media |  |  |  |  |
| --- | --- | --- | --- | --- |
| Day 5-6 |  |  |  |  |
|  | Stock conc.: |  | final conc.: |  |
| Ascorbic Acid, 0.25M stock | 250 | mM | 0.25 | mM |
| FGF-7, 500µg/mL stock | 500,000 | ng/mL | 50 | ng/mL |
| Retinoic Acid, stock 10mM | 10,000 | µM | 1 | µM |
| SANT-1, 0.25µM | 2,500 | µM | 0.25 | µM |
| TPB, 200nM final | 1,000,000 | nM | 200 | nM |
| LDN, 100nM final | 1,000,000 | nM | 100 | nM |
| ITS-X | 200 | x | 1 | x |
| IWP-6, 5µM final | 10,000 | µM | 5 | µM |

| Stage 4 Pancreatic Endoderm (EP) media |  |  |  |  |
| --- | --- | --- | --- | --- |
| Day 7-9 |  |  |  |  |
|  | Stock conc.: |  | final conc.: |  |
| Ascorbic Acid, 0.25mM | 250 | mM | 0.25 | mM |
| FGF-7, 100µg/mL (diluted stock 1:5) | 100,000 | ng/mL | 2 | ng/mL |
| Retinoic Acid, stock: 1mM (diluted) | 1,000 | µM | 0.1 | µM |
| SANT-1, 0.25µM | 2,500 | µM | 0.25 | µM |
| TPB, 100nM | 1,000,000 | nM | 100 | nM |
| LDN, 200nM | 1,000,000 | nM | 200 | nM |
| ITS-X, 1:200 dilution | 200 | x | 1 | x |

Similarly, for the final stages of the differentiation basal media were prepared in advance and each day compounds were added according to below scheme to obtain Endocrine Precursors (EP) on Day 13 and β-like cells on Day 20.

| S5-S6-S7 basal media |  |  |  |  |
| --- | --- | --- | --- | --- |
|  | Stock conc.: | stock unit | final conc.: | unit |
| Sodium bicarbonate |  |  | 1.5 | g/L |
| Glucose, 10mM, (1000mg/L already) | 200000 | g/L | 3600 | mg/L Total |
| BSA, 0.5% | 10% | BSA | 2.0% |  |
| Glutamax | 100 | x | 1 | x |

| Stage 5 Endocrine Precursor/progenitor (EP) |  |  |  |  |
| --- | --- | --- | --- | --- |
| Day 10-12 |  |  |  |  |
|  | Stock concentration |  | Final Concentration |  |
| Retinoic Acid, stock: 1mM (diluted) | 1,000 | μM | 0.05 | μM |
| SANT-1, 0.25μM | 2,500 | μM | 0.25 | μM |
| LDN, 100μM | 1,000,000 | μM | 100 | μM |
| ITS-X | 200 | x | 1 | x |
| T3, 1μM | 30 | μM | 1 | μM |
| Alk5i, 10μM | 10,000 | μM | 10 | μM |
| Zinc Sulfate, 10μM | 10,000 | μM | 10 | μM |
| Heparin, 10μg/mL | 50,000 | μg/mL | 10 | μg/mL |

| Stage 6: β-like stage |  |  |  |  |
| --- | --- | --- | --- | --- |
| Day 13-19 |  |  |  |  |
|  | Stock concentration |  | Final Concentration |  |
| LDN | 1,000,000 | nM | 100 | nM |
| ITS-X | 200 | x | 1 | x |
| T3 | 29.72 | μM | 1 | μM |
| Alk5i, 10mM | 10,000 | μM | 10 | μM |
| Zinc Sulfate, 10mM | 10,000 | μM | 10 | μM |
| Heparin, 10μg/mL | 50,000 | μg/mL | 10 | μg/mL |
| Gamma-sec inh, XX, cat. 565789, 100μM stock | 100,000 | nM | 100 | nM |

### Western blot

For western blot cells were harvested at day 13 from a 12-well dish, washed 1x in PBS then scraped off in 120µL RIPA buffer including cOmplete Ultra Protease inhibitors (Roche), lysed on ice for 10minutes and then sonicated 5×30sec ON/OFF on a Diagenode BioRuptor in 1.5mL eppendorff tubes. Subsequently spun down at 21 000g for 30 minutes at 4°C, saving the supernatant. Pierce BCA protein kit (ThermoFisher) was used to measure protein concentration on a Nanodrop 2000 (ThermoFisher). Ran 30 µg protein on a NuPage 4-12% BisTris SDS-PAGE gel in MOPS buffer (ThermoFisher) and transferred using BioRad Mini-Protean transfer system. Blocked in 5% skim milk in PBS-tween 0.1% for 1 hour at room temperature. Incubated with primary antibodies according to below scheme over-night 4°C, washed 3×10 minutes in PBS-tween 0.1%, then incubated with secondary antibodies (Jackson Labs, raised in Donkey) for 1h at room temperature then washing 3×10 minutes in PBS-tween 0.1%.

Horseradish peroxidase conjugated antibody blot were developed using ECL Prime Western Blotting System (Sigma) and captured both HRP and fluorescent label on the ChemiDoc system (BioRad).

| Primary Antibody | Dilution | Company | Cat. No. | Species | Label on Secondary ab | Dilution |
| --- | --- | --- | --- | --- | --- | --- |
| **α-HES1 (H-140)X** | 1:2000 | Santa Cruz | SC-25392X | Rabbit | HRP | 1:5000 |
| **α-VINCULIN** | 1:5000 | Sigma | V9131 | Mouse | Cy3 | 1:2500 |

Horseradish peroxidase conjugated antibody blot were developed using ECL Prime Western Blotting System (Sigma) and captured both HRP and fluorescent label on the ChemiDoc system (BioRad).

### Confocal Immunofluorescence Microscopy

Cells for immunofluorescence were grown on Ibidi µ-slides 8-well (cat. No. 80826) and washed once in PBS+/+, then fixed in 4% Formaldehyde (=10% Formaline, VWR) for 30 minutes. Then washed twice in PBS+/+. Permeabilised with 0.5% Triton X-100 in PBS for 10minutes, then washed once in PBS+/+ and blocked in SuperBlock (ThermoFisher) for 30 minutes. Primary antibodies were diluted 0.1% Triton X-100 in PBS according to below scheme and incubated overnight at 4°C, then washed 3×5 minutes in PBS. Secondary antibodies (1:500, Jackson Lab, raised in Donkey) were incubated for 45 minutes at room temperature and then washed 3x in PBS.

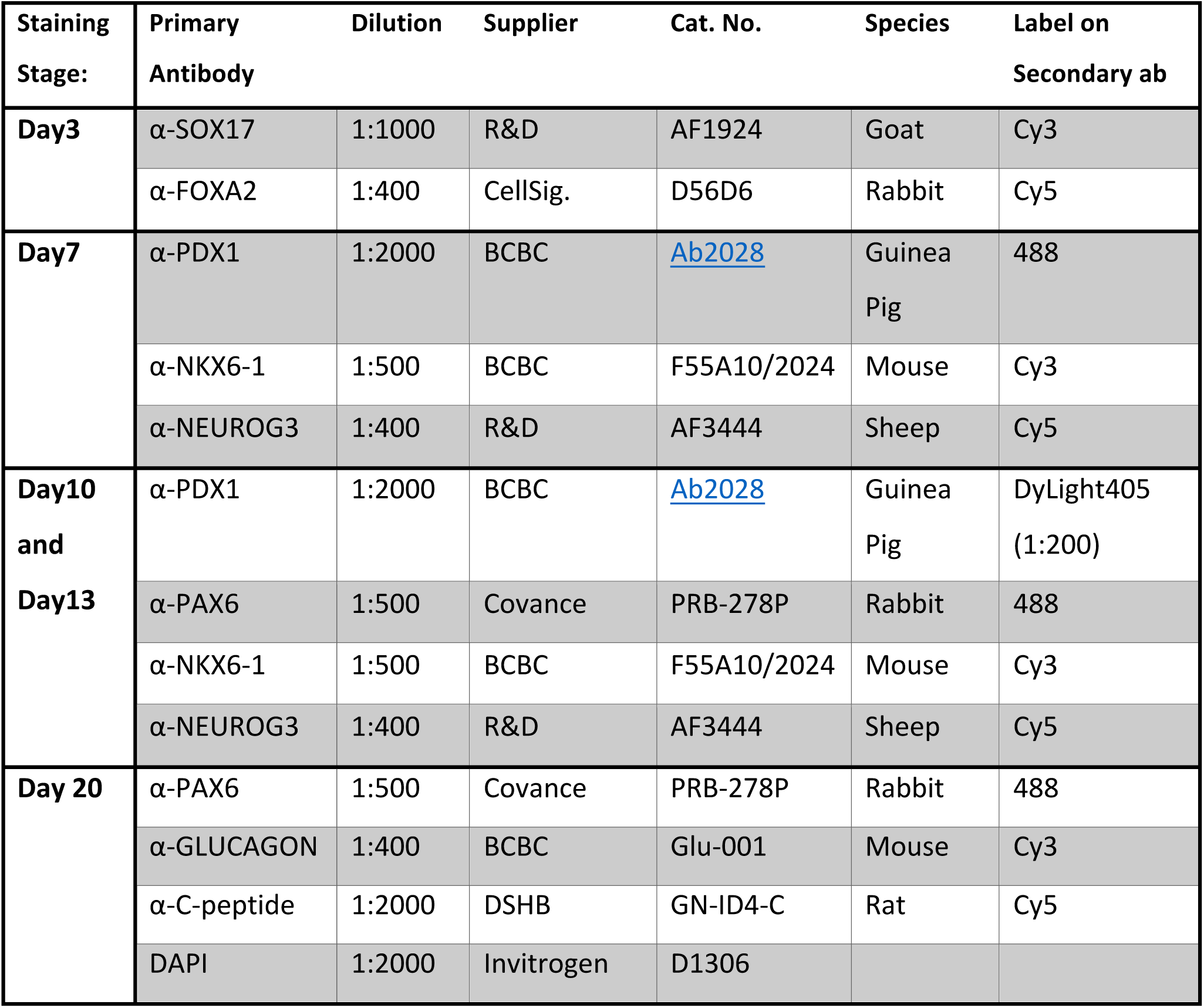

| Staining Stage: | Primary Antibody | Dilution | Supplier | Cat. No. | Species | Label on Secondary ab |
| --- | --- | --- | --- | --- | --- | --- |
| Day3 | α-SOX17 | 1:1000 | R&D | AF1924 | Goat | Cy3 |
| | α-FOXA2 | 1:400 | CellSig. | D56D6 | Rabbit | Cy5 |
| Day7 | α-PDX1 | 1:2000 | BCBC | Ab2028 | Guinea Pig | 488 |
| | α-NKX6-1 | 1:500 | BCBC | F55A10/2024 | Mouse | Cy3 |
| | α-NEUROG3 | 1:400 | R&D | AF3444 | Sheep | Cy5 |
| Day10 and Day13 | α-PDX1 | 1:2000 | BCBC | Ab2028 | Guinea Pig | DyLight405 (1:200) |
| | α-PAX6 | 1:500 | Covance | PRB-278P | Rabbit | 488 |
| | α-NKX6-1 | 1:500 | BCBC | F55A10/2024 | Mouse | Cy3 |
| | α-NEUROG3 | 1:400 | R&D | AF3444 | Sheep | Cy5 |
| Day 20 | α-PAX6 | 1:500 | Covance | PRB-278P | Rabbit | 488 |
| | α-GLUCAGON | 1:400 | BCBC | Glu-001 | Mouse | Cy3 |
| | α-C-peptide | 1:2000 | DSHB | GN-ID4-C | Rat | Cy5 |
|  | DAPI | 1:2000 | Invitrogen | D1306 |  |  |

Confocal images were captured on Leica SP8 at 1024×1024 with line average of 4 and stitched together. Adjustment of brightness and contrast was done in Adobe Photoshop with the legacy option.

### RT-qPCR

For Reverse Transcription Quantitative Polymerase Chain Reaction (RT-qPCR) cells from a 12-well were washed once in PBS and then harvested using RNeasy Mini plus kit on a Qiacube (Qiagen). Quality and quantity were measured on a Nanodrop 2000 and 1µg of total RNA was used for RT reaction with Superscript III (ThermoFisher) according to Manufacturer’s instructions using a mix of Random Hexamers and Oligo-dT primer in a 20 µL reaction. Each Sample by Gene was done in duplicate of 10µL reaction using 4µL 1:10 diluted cDNA, 1µL 5µM primer mix and 5µL Lightcycler 480 SYBR Green I MM (Roche). Run in a 384 well plate format on a Lightcycler 480 (Roche) using the standard SYBR protocol.

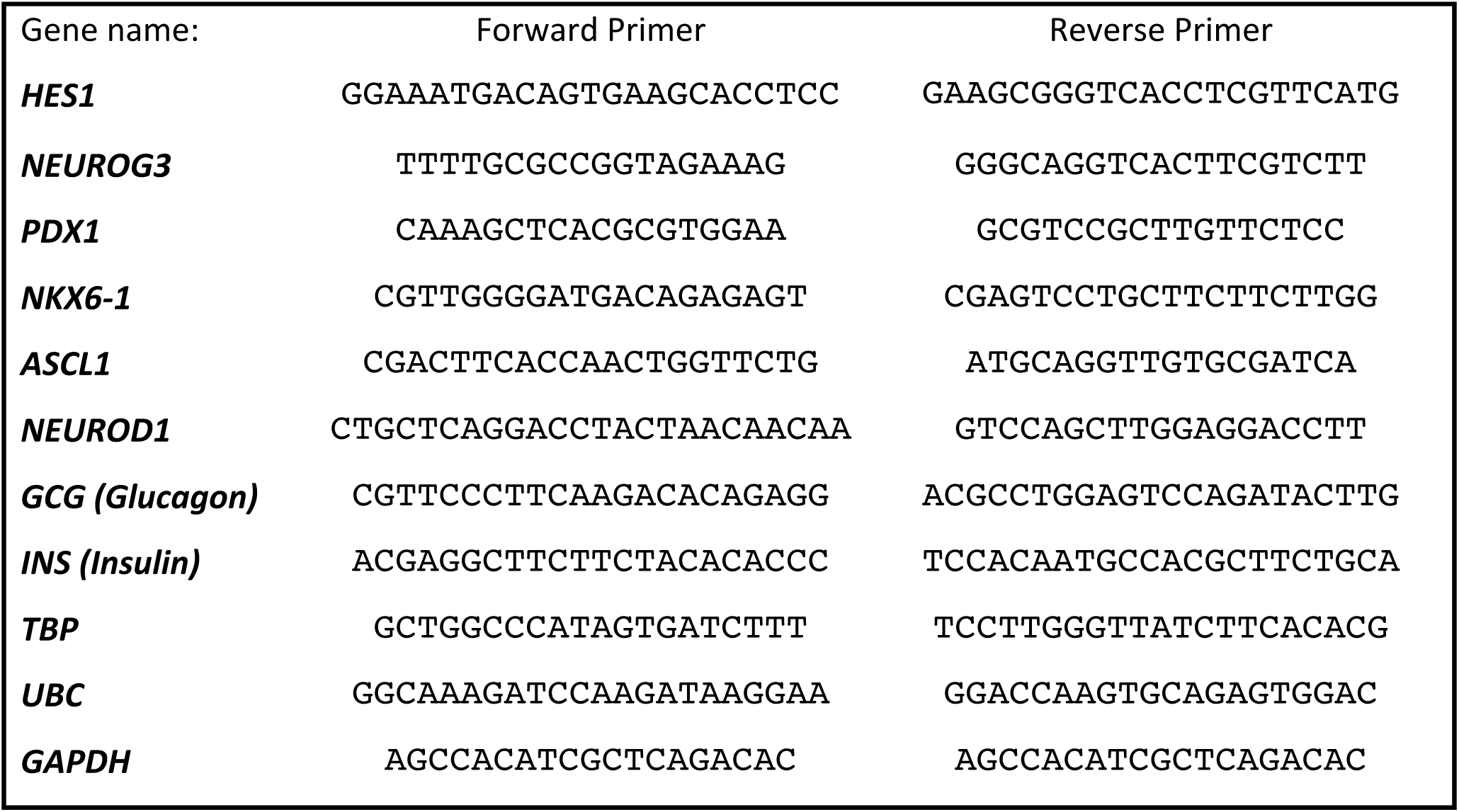

Expression values are normalised to the geometric mean of the housekeeping genes TBP, UBC and GAPDH after testing for lowest coefficient of variation.

### RNA-seq

For building RNA-seq libraries two cell lines from each genotype and from differentiation #4 and #5 with 1µg of RNA using the NEB NEXT Ultra II with mRNA magnetic isolation module (NEB #E7775 and #E7490) with 5 cycles of amplification. Quality of the RNA and subsequently of the libraries were measured on a Fragment Analyser and the libraries were loaded accordingly on Illumina NextSeq 500 with Hi-output 1×75bp kit.

| Sample No. | Sample ID | Index<br>(barcode) | Million Aligned Reads<br>(STAR) |
| --- | --- | --- | --- |
| 1 | Diff4 Day13 H1 | ATCACG | 26.62 |
| 2 | Diff4 Day13 22 | CGATGT | 27.13 |
| 3 | Diff4 Day13 21sub10 | TTAGGC | 21.66 |
| 4 | Diff4 Day13 24sub3 | TGACCA | 25.33 |
| 5 | Diff4 Day13 H3 | ACAGTG | 26.08 |
| 6 | Diff4 Day13 101 | GCCAAT | 27.61 |
| 7 | Diff4 Day13 28 | CAGATC | 25.89 |
| 8 | Diff4 Day13 32 | ACTTGA | 27.52 |
| 9 | Diff5 Day13 H1 | GATCAG | 29.42 |
| 10 | Diff5 Day13 22 | GTGAAA | 20.79 |
| 11 | Diff5 Day13 21sub10 | GTGGCC | 29.61 |
| 12 | Diff5 Day13 24sub3 | GTTTCG | 27.76 |
| 13 | Diff5 Day13 H3 | CGTACG | 28.08 |
| 14 | Diff5 Day13 101 | GAGTGG | 27.11 |
| 15 | Diff5 Day13 28 | ACTGAT | 28.15 |
| 16 | Diff5 Day13 32 | ATTCCT | 25.41 |

### RNA-seq bioinformatics analysis

Reads from the Nextseq were converted to FASTQ-files using bcl2Fastq (Illumina) on Computerome high performance cluster, were also FASTQC quality control and alignment to hg38 human genome with STAR was done(Dobin et al., 2013) with standard settings resulting in >90% alignment of reads. Loaded the gene count table into DESEQ2 for differential expression analysis on gene level (Love et al., 2014). With the DESEQ2 package in R we performed quality control assessment by Principal Component Analysis after regularized logarithmic transformation. For the differential expression analysis we did Log-Fold-Change shrinkage with DESEQ2 and tested Wildtype vs. HES1^-/-^, Wildtype vs. NEUROG3^-/-^, Wildtype vs. HES1^-/-^NEUROG3^-/-^, HES1^-/-^ vs. HES1^-/-^NEUROG3^-/-^ and NEUROG3^-/-^ vs. HES1^-/-^NEUROG3^-/-^. Altogether this resulted in 2063 genes deregulated in at least one of these comparisons, used for the graphs in (Figure 3).

Using the org.Hs.eg.db for annotation and EDAseq packages gene lengths we obtained gene length normalised read counts (FPKM), which is useful for comparing absolute expression levels across genes. For plotting ggplot was mostly except for heatmaps and kmeans clustering (centers=5) for which superheat package was used with manhattan distance measure and ward.D2 method.

Gene Set Enrichment Analysis was done with KEGG database, transcriptomics form our own lab described below and gene sets from recent publications using single cell RNA-sequencing to find markers exclusively expressed in specific cell types in order to establish a database of 42 cell types from human pancreas, mouse intestine (lifted to human) and neuronal cell types (Haber et al., 2017; La Manno et al., 2016; Segerstolpe et al., 2016). Overview of database genesets from outside sources. We also employed datasets form our own lab, including genes up-and down-regulated on RNA-seq from FACS sorted GFP+ cells from E10.5 *Hes1*^+/+^ *Pdx1*^GFP/+^ and *Hes1*^-/-^*Pdx1*^GFP/+^ mouse embryos from Manuscript III (Jørgensen et al.), and a comprehensive microarray analysis of isolated bi-potent trunk progenitors, endocrine precursors and pre-acinar cells obtained by from FACS sorting of DBA-stained, dissociated E15.5 embryonic pancreas cells from Hes1-GFP; Neurog3-tRFP embryos (K. Honnens de Lichtenberg et al., Manuscript I). Mouse data were lifted to human annotation using Homologene database through R. For GSEA analysis the DESEQ2 results were imported based on Wald Statistics as a preranked lists and enrichement was calculated with the classic setting and 1000 permutations.

| SingleCell-RNA-seq datasets for GSEA: | Cell Type | Tissue |
| --- | --- | --- |
| Segerstolpe_SC_beta | Beta cell | Adult Human Pancreas |
| Segerstolpe_SC_alpha | Alpha cell | Adult Human Pancreas |
| Segerstolpe_SC_gamma | Gamma cell | Adult Human Pancreas |
| Segerstolpe_SC_delta | Delta cell | Adult Human Pancreas |
| Segerstolpe_SC_acinar | Acinar cell | Adult Human Pancreas |
| Segerstolpe_SC_ductal | Ductal cell | Adult Human Pancreas |
| Segerstolpe_SC_epsilon | Epsilon cell | Adult Human Pancreas |
| Haber_Goblet | Goblet | Mouse Intestine |
| Haber_Paneth | Paneth | Mouse Intestine |
| Haber_Tuft | Tuft | Mouse Intestine |
| Haber_Enteroendocrine | Enteroendocrine | Mouse Intestine |
| Haber_ProxEnterocyte | Proximal Enterocyte | Mouse Intestine |
| Haber_DistEnterocyte | Distal Enterocyte | Mouse Intestine |
| Haber_Early_EnteroEndocrine_Progenitor | Enteroendocrine progenitor, Early | Mouse Intestine |
| Haber_Late_EnteroEndocrine_Progenitor | Enteroendocrine progenitor, Mid | Mouse Intestine |
| Haber_Mid_EnteroEndocrine_Progenitor | Enteroendocrine progenitor, Late | Mouse Intestine |
| Basal_LaMannoGyllborg_hBrain_scRNAseq | Basal | Human Brain |
| hEndo_LaMannoGyllborg_hBrain_scRNAseq | hEndo | Human Brain |
| hPeric_LaMannoGyllborg_hBrain_scRNAseq | hPeric | Human Brain |
| hMgl_LaMannoGyllborg_hBrain_scRNAseq | hMgl | Human Brain |
| hOPC_LaMannoGyllborg_hBrain_scRNAseq | hOPC | Human Brain |
| hRgl2c_LaMannoGyllborg_hBrain_scRNAseq | hRgl2c | Human Brain |
| hRgl2b_LaMannoGyllborg_hBrain_scRNAseq | hRgl2b | Human Brain |
| hRgl2a_LaMannoGyllborg_hBrain_scRNAseq | hRgl2a | Human Brain |
| hRgl3_LaMannoGyllborg_hBrain_scRNAseq | hRgl3 | Human Brain |
| hRgl1_LaMannoGyllborg_hBrain_scRNAseq | hRgl1 | Human Brain |
| hProgM_LaMannoGyllborg_hBrain_scRNAseq | hProgM | Human Brain |
| hProgBP_LaMannoGyllborg_hBrain_scRNAseq | hProgBP | Human Brain |
| hProgFPL_LaMannoGyllborg_hBrain_scRNAseq | hProgFPL | Human Brain |
| hProgFPM_LaMannoGyllborg_hBrain_scRNAseq | hProgFPM | Human Brain |
| hNProg_LaMannoGyllborg_hBrain_scRNAseq | hNProg | Human Brain |
| hNbM_LaMannoGyllborg_hBrain_scRNAseq | hNbM | Human Brain |
| hNbML1_LaMannoGyllborg_hBrain_scRNAseq | hNbML1 | Human Brain |
| hRN_LaMannoGyllborg_hBrain_scRNAseq | hRN | Human Brain |
| hNbML5_LaMannoGyllborg_hBrain_scRNAseq | hNbML5 | Human Brain |
| hDA0_LaMannoGyllborg_hBrain_scRNAseq | hDA0 | Human Brain |
| hDA1_LaMannoGyllborg_hBrain_scRNAseq | hDA1 | Human Brain |
| hDA2_LaMannoGyllborg_hBrain_scRNAseq | hDA2 | Human Brain |
| hNbGaba_LaMannoGyllborg_hBrain_scRNAseq | hNbGaba | Human Brain |
| hGaba_LaMannoGyllborg_hBrain_scRNAseq | hGaba | Human Brain |
| hSert_LaMannoGyllborg_hBrain_scRNAseq | hSert | Human Brain |
| hOMTN_LaMannoGyllborg_hBrain_scRNAseq | hOMTN | Human Brain |

### Chromatin Immunoprecipiation with Next Generation Sequencing

For Chromatin immunoprecipitation with Next Generation Sequencing (ChIP-seq) we used the HUES4 PDX-GFP reporter human embryonic stem cell line (kind gift from H. Semb and J. Ameri). The reporter cell line was seeded on Fibronectin at cells/cm2 and followed the differentiation protocol as described above. On Day 13 cells were dissociated with TrypLE and Accutase (ThermoFisher) mix 1:1 for 15 minutes to single cell suspension, spun down 5 minutes 500g and resuspended in FACS buffer (PBS with 10% BSA). Around ¼ was used as an unsorted sample, for the rest added DAPI for live/dead stain and set gates according to MEF GFP-population on BD ARIA II for sorting. Recovered ∼3 million PDX-GFP^+^ cells per experiment from the sorting and treated them in parallel with the unsorted as follows: spun down 500g for 5 minutes with low acceleration/deceleration and resuspended pelleted cells in 1% Formaldhyde in PBS (freshly made from Amersham MeOH-free 16% Formaldehyde) for 10 minutes. Stopped fixation in 0.125M Glycine for 5 minutes and washed twice in PBS. Then did nuclear enrichment in 0.5% SDS ChIP buffer for 5 minutes and discarded supernatant after spinning. Lysed in RIPA buffer with protease inhibitors for 10 minutes on ice. Sonicated 7×30seconds ON/OFF, then span full speed for 20 minutes. For the PDX-GFP+ samples 2.5µg chromatin per ChIP and Unsorted 10µg chromatin per ChIP was diluted to 500µL in RIPA buffer. Tumbled overnight at 4°C with primary antibodies 1µL HES1 (H140-X / SC-25392X) and 1µL rabbit IgG (#2729, Cell Signaling). Added Protein G Dynabeads and tumbled for 4 hours, washed in RIPA + inhibitors twice, then once in TE buffer moving to a fresh eppendorff tube and eluted in Elution buffer containing 1%NaHCO3, 1% SDS and proteinase K in PBS for 2h at 56°C and then overnight at 65°C. Cleaned up ChIP’ed DNA with ChIP Clean and Concentrator kit (Zymo) and quantified on a Qubit (ThermoFisher) for sequencing. Built libraries from ∼1500pg ChIP’ed DNA for multiplex libraries using a protocol developed in the Ido Amit lab (Blecher-Gonen et al., 2013) and sequenced on an Illumina HiSeq 2500 at the Weizmann Institute, Israel.

### ChIP-seq Bioinformatics analysis

FastQ files were aligned to the human hg19 genome using Bowtie2 (Langmead and Salzberg, 2012), converted the SAM-files to BAM and indexed bam files using SamTools and made bigwig tracks for visualisation using Deeptools. Tracks were loaded into Integrated Genome Viewer (IGV) 2.4 with HES1 tracks, peaks and motif-search as indicated in figures.

Used the two replicates of unsorted HES1 ChIP for MACS2 peak calling since they were of superior quality. After close evaluation of peaks called in HES1-ChIP #1 (160209) the top 4000 most enriched were used, resulting in 3750 peaks after removing areas on the UCSC “blacklist”. For HES1-ChIP #2 (160322) which is less enriched we set a relaxed p-value cutoff of 0.002 to yield 4490 peaks. The two peaklists were overlapped using bedtools for the 998 high confidence HES1 peaks and many were checked manually for verification of thresholds. For defining transcription start site (TSS) the hg19 TSS were downloaded from the table browser of known UCSC genes at genome.UCSC.edu. Used summit-peak file +200 bp for Homer de Novo motif search from either the peak-set overlapping TSS or not overlapping the TSS. From previously published data from the Ferrer and Hattersley labs FOXA2 from a similar stage of human ES differentiation (E-MTAB-1990)(Weedon et al., 2014). Other published dataset were also in consideration including histone marks such as H3K4me3, H3K27me3 from the Sander lab but did not have proper comparative quality (Xie et al., 2013).

Peak list was annotated with GREAT (McLean et al., 2010) using basal+extension settings, including curated regulatory domains to yield 1049 genes associated with HES1 peaks.

For overlap of HES1 ChIP-seq peaks and RNA-seq expression data the gene name was used, recognising all the HES1 associated genes.

